# *Plasmodiophora brassicae* chitin-binding effectors guard and mask spores during infection

**DOI:** 10.1101/2020.12.23.423615

**Authors:** Kevin Muirhead, Edel Pérez-López

## Abstract

Plants have a sophisticated and multilayered immune system. However, plant pathogens, helped by effector proteins, have found several strategies to evade plant immunity. For instance, the clubroot pathogen, *Plasmodiophora brassicae,* is able to turn the roots of the susceptible hosts into nutrient-sink galls surpassing patterns-triggered immunity (PTI) and effector-triggered immunity (ETI). Chitin, the main component of *P. brassicae* spores cell walls and a well-known pathogens-associated molecular pattern (PAMP), can elicit PTI but is also the target of plant chitinases and chitin deacetylases. The fact that *P. brassicae* does not trigger PTI during the infection of the susceptible hosts motivated a genome-wide search of genes coding for secreted chitin-related proteins. We found that *P. brassicae* genome encodes a large repertoire of candidate-secreted effectors containing the chitin-binding domain carbohydrate-binding module family 18 (CBM18), along with chitinases and chitin deacetylases domains. The role of such proteins in the pathogenicity of the clubroot pathogen is unknown. Here, we characterized the function of two effectors, *Pb*ChiB2 and *Pb*ChiB4, which are transcriptionally activated during the spores transition to uninucleate primary plasmodium and during the spore formation. Through co-precipitation, we found that recombinant *Pb*ChiB2 and *Pb*ChiB4 bind to the spores and to chitin oligomers *in vitro*. We also showed that both proteins suppress chitin-triggered activation of the immune MPK3 and MPK6 in the host *Brassica napus*. These findings suggest a dual role for the *P. brassicae* CBM18 proteins as effectors for protecting zoospores and resting spores formation and for suppressing chitin-triggered immunity during the infection.

## INTRODUCTION

Pathogen colonization is an essential step of infection extremely guarded by the plant hosts (Zipfel 2014). Through pattern-recognition receptors (PRRs) located on the cell surface, plants can perceive pathogen-associated molecular patterns (PAMPs) and the activation of PRR-triggered immunity (PTI) takes place (Dodds and Rathjen 2010). This first layer of immunity is characterized by the generation of reactive oxygen species, the activation of channels, and the phosphorylation of mitogen-activated protein kinases leading to a wide array of antimicrobial defense mechanisms (Dodds and Rathjen 2010; Zipfel 2014). However, plant pathogens have developed very sophisticated strategies to scape PTI, infect the plant host, and to complete their life cycle (Dodds and Rathjen 2010).

Among other very well studied PAMPs, chitin [polymer of b-1,4-linked N-acetylglucosamine (GlcNAc)n] has been getting a lot of attention by the MPMI community (Volk et al. 2019; Cheval et al. 2020). Chitin is a major component of fungal cell walls, and it has been shown that during the infection, chitin oligosaccharides resulting from the activity of plant chitinases, are recognized by receptor kinases activating PTI (Sánchez-Vallet et al. 2015). To escape chitin-triggered immunity, fungal pathogens have developed several strategies including the secretion of chitin-binding effectors (van den Burg et al. 2006). These effectors can act, not only capturing free chitin oligosaccharides, but also as a shield to protect the pathogens from plant chitinases (van Esse et al. 2007; Marshall et al. 2011). The chitin-binding domains identified in fungal plant pathogen effectors can be divided into three major groups based on their carbohydrate-binding module (CBM) family: (*i*) CBM family 14 (CBM14), identified in chitin-binding effectors from *Cladosporium fulvim* (van den Burg et al. 2006) and *Mycosphaerella fijiensis* (Stergiopoulos et al. 2010); (*ii*) CBM family 50 (CBM50), also known as LysM effectors, identified in *Cladosporium fulvim* (de Jonge et al. 2010), *Zymoseptoria tritici* (Marshall et al. 2011), *Magnaporthe oryzae* (Mentlak et al. 2012), *Colletotrichum higginsianum* (Takahara et al. 2016), and *Rhizoctonia solani* (Dölfors et al. 2019); and (*iii*) CBM family 18 (CBM18), identified in *Magnaporthe oryzae* (Mochizuki et al. 2011) and *Verticillium nonalfalfae* (Volk et al. 2019). While chitin is predominantly found in cell walls of fungal plant pathogens, it is also the main carbohydrate present in resting spore cell walls of in *Plasmodiophora brassicae* Woronin, the clubroot pathogen.

Clubroot is a devastating disease affecting cruciferous crops worldwide. It is caused by the obligate parasite, biotrophic, soil-borne, parasitic protist *P. brassicae* (Burki et al. 2010). The life cycle of *P. brassicae* takes place in the roots of the infected host. It is divided in two stages: (*i*) primary infection, characterized by the encysting of the spores, penetration through the root hairs, and the subsequent formation of the zygote, and (*ii*) secondary infection, stage of the life cycle characterized by the formation of the secondary plasmodium and finally release back to the soil of the resting spores (Liu et al. 2020). Although *P. brassicae* secondary infection has drawn the most attention because it leads to the host death, primary infection remains the key step for infection (Pérez-López et al. 2020). During the analysis of *P. brassicae* transcriptome in different hosts and developmental stages, several proteins with the chitin-binding (CBM18) domain were enriched in *P. brassicae* secretome (Schwelm et al. 2015), although the function of these proteins remains unknown. Similar results were obtained in another study looking into effector proteins involved in *P. brassicae* primary infection (Chen et al. 2019).

Here we performed a genome-wide search to identify all the chitin-related proteins putatively secreted by *P. brassicae* with special focus on those bearing CBM18 domains. Among the proteins identified, we characterized *Pb*ChiB2 and *Pb*ChiB4 which have two CBM18 modules and a functional signal peptide. Both are presumably secreted during resting spores development to plasmodium and during resting spores formation. We show that both proteins bind resting spores and chitin *in vitro* and that these proteins are able to inhibit chitin-triggered immunity ‘guarding’ the resting spores throughout the infection.

## MATERIALS AND METHODS

### Plant material and growth conditions

*Arabidopsis thaliana* Col-0 and rapeseed (*B. napus,* canola in Canada) Westar seeds, disinfected with 70% ethanol, 95% ethanol, 8% bleach and sterile water, were sown on 1/2 Murashige and Skoog (MS) salts agar (1%, w/v) plates (Sigma-Aldrich, CAD) with 1% sucrose. Plates were placed at 4 °C for 4 days and then transferred to a growth chamber (Conviron ATC10, CMP4030 control system; 170 μmol photons m^2^ s^−1^; 16 h/8 h light/dark cycle; 20 °C) for germination and growth.

### Clubroot infection

Ten-day old Arabidopsis seedlings were transferred to Vèrada soil mix (Farad, CAD) with four plants in each ‘square pot’ (3.5 × 3.5 inch). Seedlings were allowed to acclimatise in these pots for four additional days before 500 uL of 5×10^7^ resting spores mL^−1^, extracted from canola galls (Pérez-López et al. 2020) was applied to each plant where the stem entered the soil. Control plants were inoculated with 500 μL of distilled water (mock inoculation) and grown in separate trays in the same growth chamber.

### Bioinformatics analyses

To identify putative secreted chitin-related proteins we performed a genome-wide search for proteins having a signal peptide prediction score ≥ 0.7, and a length of < 500 amino acids in the *P. brassicae* Pbe3.h15 genome (Schwelm et al. 2015). The resulting proteins were analyzed using Pfam (https://pfam.xfam.org/) to identify chitin-related Carbohydrate-Active enZymes (chitin-CAZymes). The proteins identified were characterized using several *online* tools as previously described (Pérez-López et al. 2020), including ApoplastP (Sperschneider et al. 2017) to know their putative apoplastic localization. The expression of the genes coding these proteins was analyzed using the RNAseq data previously generated by (Schwelm et al. 2015) under the SRA accession numbers ERX1409397 - ERX1409403. RNAseq libraries were downloaded using the fastq-dump program of the NCBI SRA Toolkit v 2.9.6 (Leinonen et al. 2011). The RNAseq libraries were assessed for quality using fastqc v 0.11.8 (Andrews 2010) and TruSeq3 Illumina adapters were determined from the most represented sequences. Reads were quality-filtered and trimmed using trimmomatic v 0.39 (Bolger et al. 2014) with the following parameter settings ILLUMINACLIP:TruSeq3-PE.fa:2:30:10:2:keepBothReads LEADING:3 TRAILING:3 MINLEN:36. Reads were mapped to the P. brassicae Pbe3.h15 genome (Schwelm et al. 2015) using STAR v 2.7.5c (Dobin et al. 2013). The STAR genome database was generated using the following parameter settings --genomeSAindexNbases 11 --runMode genomeGenerate. STAR was executed using the following parameter settings --runMode alignReads --outSAMtype SAM --outSAMattributes All --outSAMstrandField intronMotif. The SAM output files from STAR were processed using samtools v 1.9 and analyzed using the cuffdiff program of Cufflinks v2.1.1 (Trapnell et al. 2010). The heatmap was generated using cummeRbund 3.12 (Goff et al. 2020) package using the output generated by the cuffdiff program and gene ids shown in Table 1. The phylogenetic analysis of chitin-CAZymes was carried out using MUSCLE v3 (Edgar 2004) and MEGA v6 (Tamura et al. 2013). The phylogenetic tree was constructed using maximum-likelihood, the James– Taylor–Thorthon model and a bootstrap value of 1000.

**Table 1.** Description and identity of the putative secreted chitin-CAZymes reported in this study.

| Name | Gene ID | Accession number | No aa | MW (kDa) | PI | SP cleavage | TM domain (Y/N) | Apoplastic [Y/N (score)] |
| --- | --- | --- | --- | --- | --- | --- | --- | --- |
| *PbChi1* | PBRA_005464 | CEO96860 | 477 | 50.3 | 4.9 | 21-22 | N | Y (0.8) |
| *PbChi2* | PBRA_005765 | CEO97161 | 357 | 37.9 | 6.3 | 18-19 | N | Y (0.63) |
| *PbChi3* | PBRA_003743 | CEO94930 | 400 | 44.3 | 5.8 | 23-24 | N | N (0.66) |
| *PbChiB1* | PBRA_003759 | CEO94946 | 222 | 22.9 | 7.6 | 22-23 | N | Y (0.72) |
| *PbChiB2* | PBRA_001907 | CEP01301 | 137 | 14.4 | 5.3 | 22-23 | N | Y (0.77) |
| *PbChiB3* | PBRA_002543 | CEP02278 | 336 | 35.5 | 7.7 | 20-21 | N | N (0.54) |
| *PbChiB4* | PBRA_002958 | CEP03198 | 172 | 17.9 | 4.2 | 17-18 | N | Y (0.81) |
| *PbChiD1* | PBRA_007091 | CEO98977 | 314 | 34.1 | 6.7 | 21-22 | N | N (0.54) |
| *PbChiD2* | PBRA_002230 | CEP01965 | 263 | 29.6 | 5.6 | 18-19 | N | N (0.71) |
| *PbChiD3* | PBRA_008962 | CEP02378 | 328 | 35.3 | 4.5 | 21-22 | N | N (0.59) |
| *PbChiBD1* | PBRA_003161 | CEP03401 | 387 | 40.8 | 4.8 | 29-30 | N | Y (0.84) |
| *PbChiBD2* | PBRA_005081 | CEO96410 | 372 | 40.3 | 4.7 | 18-19 | N | Y (0.59) |
| *PbChiBD3* | PBRA_002551 | CEP02584 | 355 | 37.7 | 4.2 | 21-22 | N | Y (0.8) |
| *PbChiBD4* | PBRA_007776 | CEP00042 | 402 | 43.5 | 6.8 | 18-19 | N | N (0.61) |
| *PbChiBD5* | PBRA_001295 | CEO99389 | 302 | 32.3 | 4.9 | 32-33 | N | Y (0.91) |
| *PbChiBD6* | PBRA_008942 | CEP02358 | 367 | 39 | 4.9 | 20-21 | N | Y (0.77) |
| *PbChiBT1* | PBRA_004239 | CEO95513 | 440 | 48.9 | 8.6 | 21-22 | N | N (0.73) |

### Signal peptide validation assay

To validate *Pb*ChiB1, *Pb*ChiB2, and *Pb*ChiB4 signal peptide (22, 22, and 17 aa, respectively), the 66, 66 and 51 bp fragments were amplified by PCR using as template the cDNA obtained from infected Arabidopsis plants with designated primer pairs introducing *EcoR*I at the N-terminal and *Xho*I (NEB, CAD) at the C-terminal end (Table S1). The amplified signal peptide fragments were introduced into the plasmid pSUC2 (Prof. Sophien Kamoun, The Sainsbury Laboratory) using T4 DNA ligase (Promega, USA). The yeast invertase assay was used to validate the signal peptides (Jacobs et al. 1997), following the protocol as previously described (Pérez-López et al. 2020), using the *Saccharomyces cerevisiae* yeast strain YTK12 (Dr. Hossein Borhan, AAFC-Saskatoon Research Centre). After selection in CMD-W media, we analyzed the invertase enzymatic activity as previously described (Yu et al. 2019).

### Gene expression analysis

Total RNA was extracted (Chomczynski and Mackey 1995) from infected plants at 0, 2, 5, 7, 14, 21 and 28 days post inoculation (dpi) and 2 μg was used to synthesize cDNA using the QuantiTect® Reverse Transcription Kit (Qiagen, CAD) following the manufacturer’s recommendations. Real-time qPCR was performed using iQ™ SYBR® Green Supermix (Bio-Rad, CAD) in a 20 μL final volume containing 300 nM of each primer and 2 μL of cDNA diluted 1:5 (v:v) in RNase-free water. Amplification was carried out using a magnetic induction cycler Real-Time System (Bio molecular systems, AUS), and reactions were quantified using micPCR software (v2.10.0). Each amplification used three technical replicates, results that were averaged to give the value for a single biological replicate. Results are expressed as LOG2 expression relative to *P. brassicae ELONGATION FACTOR-LIKE* (PBRA_001540) and *A. thaliana* ACTIN2 (At3g18780) expression using the comparative quantification method as previously described (Warton et al. 2004). Primers used are presented in Table S1.

### Production of recombinant proteins

For protein expression, the cDNA for *Pb*ChiB2 (Fig. S1) and *Pb*ChiB4 (Fig. S1) without the signal peptide was synthesized and cloned into plasmid pET-14b (GenScript, USA). Plasmids were used to transform *E. coli* SHuffle T7 cells (NEB, CAD). *E. coli* transformants were grown in 50 mL LB medium supplemented with ampicillin (100 μg mL^−1^) at 37 °C, 200 rpm for 3 h. Protein expression was induced by the addition of 1 mM IPTG and growth was continued at 37 °C, 200 rpm for 2 h. After centrifugation at 3200 *g* for 15 min, cell pellets were lysed using a Branson sonicator equipped with a microtip (Branson ultrasonics, USA). Pellets containing His-*Pb*ChiB2, and His-*Pb*ChiB4 were purified with Ni-NTA agarose (Thermo Fisher Scientific, CAD) through native conditions following the manufacturer‘s recommendations. Protein concentration was measured using a Qubit protein assay kit (Thermo Fisher Scientific, CAD).

### Western blot analysis

Proteins were separated by 15% SDS-PAGE and transferred to 0.2 μm PVDF membranes (Bio-Rad, CAD) for 1 h at 0.4 V cm^−1^ in a Mini-PROTEAN® Tetra Cell (Bio-Rad, CAD). His-tagged proteins were detected using monoclonal anti-polyHis-HRP conjugate (Sigma-Aldrich, CAD). Phosphorylated mitogen-activated protein kinases (MAPK) were detected using phospho-p44/p42 MAPK primary antibody (Cell Signally Technology, Leiden, Netherland), followed by a goat anti-rabbit secondary antibody HRP conjugate (Sigma-Aldrich, CAD) following the manufacturer’s recommendations. Membranes were blocked with 5% skim-milk and HPR activity was detected using SuperSignal™ West Pico PLUS Chemiluminescent Substrate (Thermo Fisher Scientific, CAD) with an Azure imager C300 (Azure Biosystems, CAD), to detect the chemiluminiscent signal.

### Carbohydrate sedimentation assay

This assay was performed as described by Volk et al. (2019). Briefly, 20 μg of recombinant *Pb*ChiB2 and *Pb*ChiB4 resuspended in 20 mM Tris (pH 8.0) were independently mixed with 2 mg of chitin magnetic beads (NEB, CAD), chitin from shrimp shell (Sigma-Aldrich, CAD), chitosan from shrimp shells (Sigma-Aldrich, CAD), cellulose (Sigma-Aldrich, CAD), or xylan (Thermo Fisher Scientific, CAD) and incubated at room temperature for 2 h with shaking at 300 rpm. After centrifuging 5 minutes at 13,000 × g, the supernatant was precipitated with cold acetone and resuspended in 40 μl of 2x SDS-PAGE buffer. The pellet was washed with 800 μL of 20 mM Tris pH 8.0 and resuspended in 40 μl Laemmli sample buffer with 13% β-mercaptoethanol. The presence of *Pb*ChiB2 and *Pb*ChiB4 was detected using western blot as described above.

A similar protocol was followed to study the ability of *Pb*ChiB2 and *Pb*ChiB4 to bind resting spores. Each recombinant protein (20 μg) was incubated with 1×10^15^ resting spores mL^−1^ extracted from canola galls. In parallel, xylem sap from canola clubs at 21 dpi, extracted as previously described (Kehr et al. 2005), was used to study the effect of chitinases on the interaction of *Pb*ChiBs with resting spores. For that, we incubated the proteins with spores preincubated with xylem sap, but also we added the xylem sap to spores preincubated with the recombinant proteins. After each treatment, the spores were pelleted, washed with 20 mM Tris pH 8.0, and the presence of *Pb*ChiB2 and *Pb*ChiB4 was analyzed as described above. All the sedimentation assays were repeated twice with identical results.

### MAPK activation assay

*Brassica napus* cultivar Westar was grown for 2 weeks in 1/2-MS liquid medium after germination and seedlings were incubated independently with (*i*) water (mock), (*ii*) 1 μM chitin [(GlcNAc)6] (Santa Cruz Biotechnology, USA), (*iii*) chitin pre-incubated with recombinant *Pb*ChiB2 or *Pb*ChiB4 (10 μM) for 1 h at room temperature, and (*iv*) *Pb*ChiB2 or *Pb*ChiB4 (10 μM) (Fig. S2). Phosphorylated MAPK3 and MAPK6 were detected as previously described (Tsuda et al. 2009) after 4 minutes of treating the seedlings. This was repeated twice with the same results.

## RESULTS

### Chitin-related *P. brassicae* CAZymes secreted repertoire

Sixteen chitin-CAZymes were identified in *P. brassicae* genome under the parameters used in this study (Table 1, Fig. 1A). These proteins were divided into four main categories: chitinases (*Pb*Chi), chitin-binding (*Pb*ChiB), chitin deacetylases (*Pb*ChiD), and chitin-binding domains (*Pb*ChiBD). An additional candidate with a chitin-binding domain (*Pb*ChiBT), tyrosinase, was also found.

**Fig. 1.**
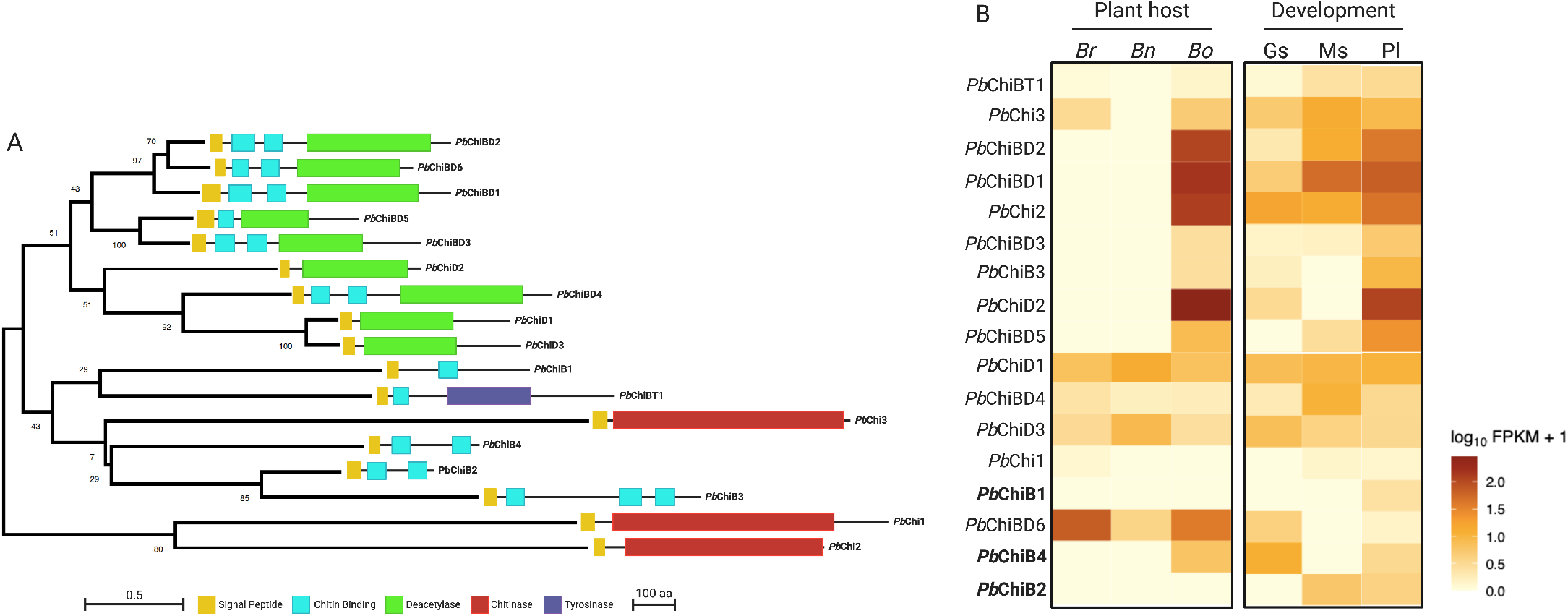
Phylogeny, structure and expression of *Plasmodiophora brassicae* chitin-CAZymes reported in this study. **A**. Phylogeny of the amino acid sequence of the chitin-CAZymes reported here through the using maximum-likelihood tree with MEGA6. The protein domain organization is on the right side of the tree. **B**. Expression profile of chitin-CAZymes measured using RNA sequencing from Schwelm et al. (2015). Overexpressed (dark orange) or underexpressed transcripts (light yellow) are presented as log FPKM+1 fold-changes relative to the mean expression measured for three *P. brassicae* hosts: *B. rapa* (Br), *B. napus* (Bn), and *B. oleracea* var. *capitate* (Bo) during secondary infection (35 dpi), and life-stage specific RNAseq data generated from germinating resting spores (Gs), maturing resting spores (Ms) and plasmodia (Pl). RNAseq data analysis scripts and output data can be found at https://github.com/kevmu/RNAseqDataSchwelm2015. Created with BioRender.com.

The domain present in the *Pb*Chi proteins was identified as a glycosil hydrolase, family 18 (GH18). The three *Pb*Chi proteins belong to the GH18 cluster A as previously reported (Schwelm et al. 2015). These proteins have less than 500 aa, a predicted signal peptide, and no transmembrane domain (Table 1). The nine *Pb*ChiD and *Pb*ChiBD were identified as members of the carbohydrate esterase family 4 (CE4) family. This family of esterases catalyze the de-acylation of polysaccharides and in all the proteins a NodB-like domain, characteristic of this family, was identified in all nine proteins (Fig. 1A). Proteins of this family have been identified as chitin deacetylases. These nine proteins have a signal peptide and four were identified as putative apoplastic proteins (Table 1). In six of the nine proteins, in addition to the NodB-like domain, we also found a chitin-binding domain (*Pb*ChiBDs) (Table 1), the carbohydrate-binding module family 18 (CBM18), with five proteins containing two copies of the domain (*Pb*ChiBD1-*Pb*ChiBD4, *Pb*ChiBD6) and *Pb*ChiBD5 with only one (Fig. 1A). Another proteins with the CBM18 domain was a tyrosinase (*Pb*ChiBT1) (Fig. 1A, Table 1).

The last four chitin-related *P. brassicae* proteins detected in this study were proteins with only the CBM18 domain (*Pb*ChiBs) (Asenio et al. 2000) (Fig. 1, Table 1). *Pb*ChiB1, with only one module of the CBM18 domain, while *Pb*ChiB2, and *Pb*ChiB4 have two, and *Pb*ChiB3 have two three modules of the CBM18 domain (Fig. 1A). Interestingly, *Pb*ChiBs with one and two modules of the CBM18 domain are predicted to be apoplastic, a common feature of chitin-binding effectors (Volk et al. 2019) (Table 1), while *Pb*ChiB3 do not seems to be apoplastic (Table 1).

### Chitin-related -encoding genes are differentially expressed in planta

To study the expression of the chitin-CAZymes and the chitin-binding proteins we used RNAseq data obtained from *P. brassicae* infected *B. rapa*, *B. napus*, and *B. oleracea* var. *capitata* during secondary infection (35 dpi), and life-stage specific RNAseq data generated from germinating resting spores, maturing resting spores and plasmodia (Schwelm et al. 2015) (Fig. 1B). In general, we observed a differential expression for each gene based on host and life-stage. Transcripts of seven of the proteins were in the same range in *B. rapa* and *B. napus* (Fig. 1B), while only one presented the same transcription profile in the three hosts (*Pb*ChiB2) (Fig. 1B). The remaining ten proteins showed a differential transcription for each host analyzed and for each life-stage. It was interesting to find that the transcript of proteins like *Pb*ChiB4 was preferentially detected in germinating spores and plasmodia but not in maturing spores (Fig. 1B), while the transcripts of other proteins like *Pb*ChiB2 were mostly detected in maturing spores and plasmodia but not in germinating spores (Fig. 1B).

### *Pb*ChiB proteins have a functional signal peptide

Moving forward, we focused on *Pb*ChiBs proteins with one and two modules of the CBM18 domain: *Pb*ChiB1, *Pb*ChiB2, and *Pb*ChiB4, in order to study their role and activity in *P. brassicae* pathogenicity. The first step was to confirm that these proteins have a functional signal peptide, indicating that they might be secreted to the apoplast. For that, the secretory activity directed by *Pb*ChiB1, *Pb*ChiB2, and *Pb*ChiB4 signal peptides was tested individually. The three signal peptides showed secretory activity in yeast as shown on CMD-W plates, and by the reduction of 2,3,5-triphenyltetrazolium chloride (TTC) to insoluble red-colored 1,3,5-triphenylformazan (TPF) (Fig. 2).

**Fig. 2.**
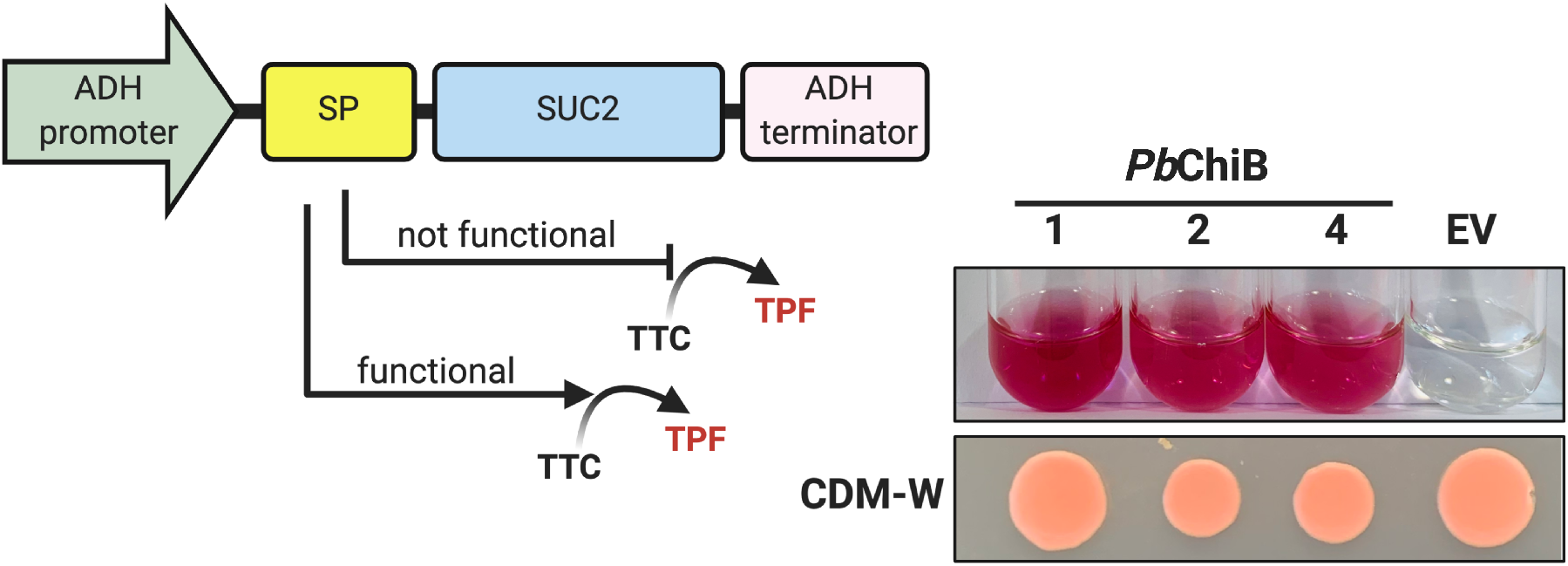
Functional analysis of *Pb*ChiB1, *Pb*ChiB2, and *Pb*ChiB4 signal peptide through yeast invertase assay. Schematic representation of the constructs containing the signal peptide sequence of *Pb*ChiB proteins and empty vector (EV) pSUC2 in yeast YTK12 were plated on CMD-W (Trp depleted), and the enzymatic activity of secreted invertase was determined using the reduction of TTC to the red-colored formazan assay. Images are representative of three replicates using independent biological samples. Created with BioRender.com.

### *Pb*ChiB2 and *Pb*ChiB4 bind chitin *in vitro*

After analyzing the chitin-binding domain present in the three *Pb*ChiB proteins, we noticed that one of the glycine of the consensus sequence was substituted by alanine in both *Pb*ChiB4 CBM18 β-hairpin loops (Fig. S3). In *Pb*ChiB1 and *Pb*ChiB2 CBM18 domains, we observed the insertion of an alanine and leucine, respectively in the consensus sequence (Fig. S3). In none of the proteins the fourth disulfide bond of the expected conserved four-disulfide core was present (Fig. S3). Based on this analysis and the fact that *Pb*ChiB1 has only one CBM18 module and weak transcripts in the three hosts analyzed and the different life-stages (Fig. 1B), only *Pb*ChiB2 and *Pb*ChiB4 were retained for further analyses (Fig. 3A).

**Fig. 3.**
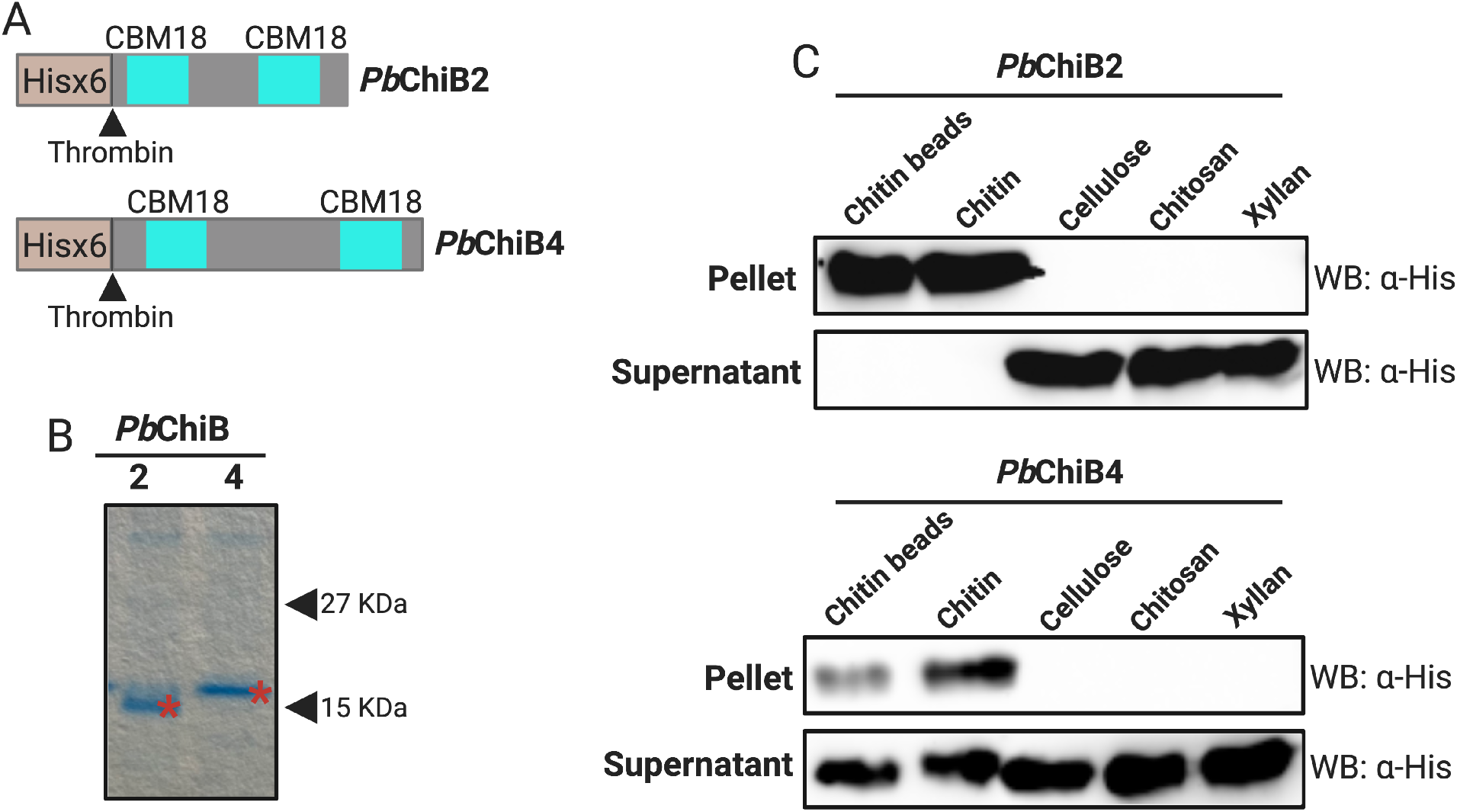
*Plasmodiophora brassicae Pb*ChB2 and *Pb*ChB4 bind specifically to chitin and chitin oligomers. **A**. Scheme of the recombinant *Pb*ChB2 and *Pb*ChB4. **B**. SDS-PAGE 15% electrophoresis of the recombinant and purified *Pb*ChB2 and *Pb*ChB4 (red star). **C**. Affinity precipitation of *Pb*ChB2 and *Pb*ChB4 in the presence of: chitin beads, shrimp shell chitin, shrimp shell chitosan, cellulose and xylan. The supernatant (S) and the pellet (P) samples were analysed after centrifugation. *Pb*ChB2 and *Pb*ChB4 was detected in the pellet fraction of chitin only using anti-His tag western blot. Created with BioRender.com.

To study the interaction with carbohydrates, recombinant and purified *Pb*ChiB2 and *Pb*ChiB4 (Fig. 3B), were used in a sedimentation assay with an array of carbohydrates. Both proteins were able to bind to chitin immobilized in magnetic beads and shrimp shell chitin, but not to shrimp shell chitosan or to the plant cell wall cellulose or xylan (Fig. 3C). Curiously, although we did not study the affinity of the interactions, we were able to detect in the pellet and the supernatant *Pb*ChiB4 while *Pb*ChiB2 was only detected in the pellet, pointing to a difference in affinity between the proteins (Fig. 3C).

### *Pb*ChiB2 and *Pb*ChiB4 are preferentially transcribed during resting spore formation

Although the transcription profile of *Pb*ChiB2 and *Pb*ChiB4 during the infection has been described (Fig. 1B), the primary infection and secondary infection consist in many different steps from 0 to 28 dpi (Liu et al. 2020). To have a better understanding of the role of these proteins, we studied their expression at four time points of primary infection (0, 2, 5, and 7 dpi), and three time points of secondary infection (14, 21, and 28 dpi) using RT-qPCR (Fig. 4A). The expression of both was initiated at 21 dpi, with a stronger induction for *Pb*ChiB2 (Fig. 4A). In addition, *Pb*ChiB4 was also strongly induced at 2dpi (Fig. 4A). Strikingly, both were strongly induced during the transition from resting sporangial plasmodium to resting spore formation (Fig. 4B), while *Pb*ChiB4 was also highly induced around primary plasmodium formation early after resting spore penetration (Fig. 4B).

**Fig. 4.**
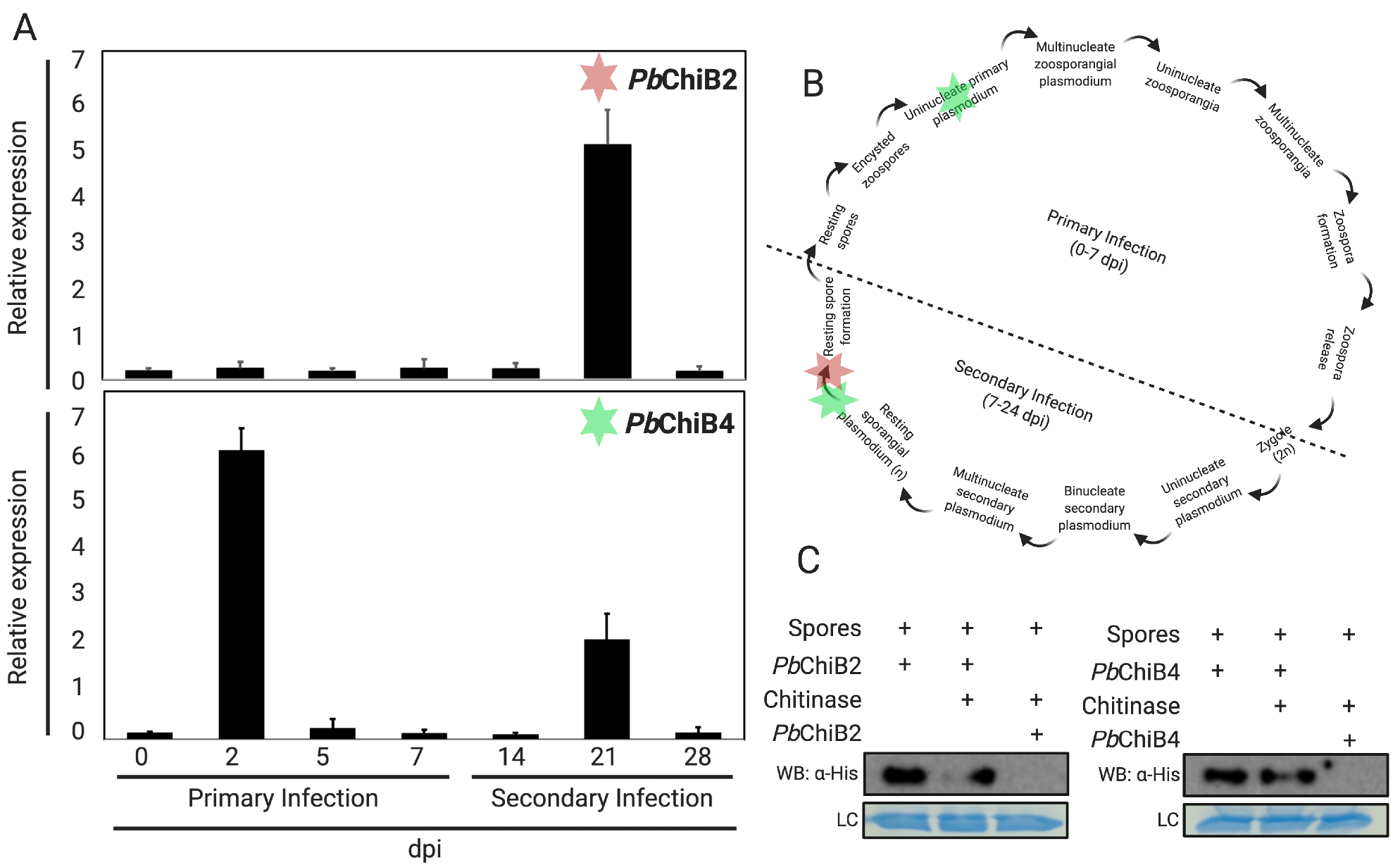
*Plasmodiophora brassicae Pb*ChB2 and *Pb*ChB4 expression profile and binding to resting spores. **A**. Expression profiling of *Pb*ChB2 and *Pb*ChB4 genes using RT-qPCR and expression levels are shown relative to the mean expression of relative to *P. brassicae ELONGATION FACTOR-LIKE* and *A. thaliana* ACTIN2. **B**. Representation of *P. brassicae* life cycle based on Liu et al. (2020) marking where *Pb*ChB2 and *Pb*ChB4 were overexpressed. **C**. Affinity precipitation of *Pb*ChB2 and *Pb*ChB4 in the presence of *P. brassicae* resting spores. The affinity was abolished when the spores were pre-incubated with xylem sap of clubroot galls (represented in the figures as Chitinases). LC loading control. Created with BioRender.com.

### *Pb*ChiB2 and *Pb*ChiB4 bind resting spores *in vitro*

To study the interaction with resting spores, recombinant *Pb*ChiB2 and *Pb*ChiB4 were used in a sedimentation assay. When *Pb*ChiB2 and *Pb*ChiB4 were incubated with resting spores isolated from canola clubroot galls, we were able to detect the His-tagged proteins in the pellet (Fig. 4C). Similar results, albeit less intense, were obtained with xylem sap from canola clubroot galls (Fig. 4C). Curiously, the interaction was abolished when the resting spores were first pre-incubated with the xylem sap before addition of the recombinant proteins (Fig. 4C).

### *Pb*ChiB2 and *Pb*ChiB4 suppress chitin-binding immunity

To study if *Pb*ChiB2 and *Pb*ChiB4 were able to compete with plant chitin receptors and interfere with early chitin-triggered immunity, we treated *B. napus* seedlings with 1μM chitin [(GlcNAc)6] and registered the activation of MAPK3 and MAPK6 4 min after application (Fig. 5A). When *Pb*ChiB2 and *Pb*ChiB4 were pre-incubated with chitin, MAPK activation was clearly reduced in the former, and to a in a lesser extent in the latter (Fig. 5B). The results obtained with *Pb*ChiB4 are in agreement with what we observed during the sedimentation assay. After treatment with water or with the *Pb*ChiB proteins alone did not induce detectable MAPK activation (Fig. 5B).

**Fig. 5.**
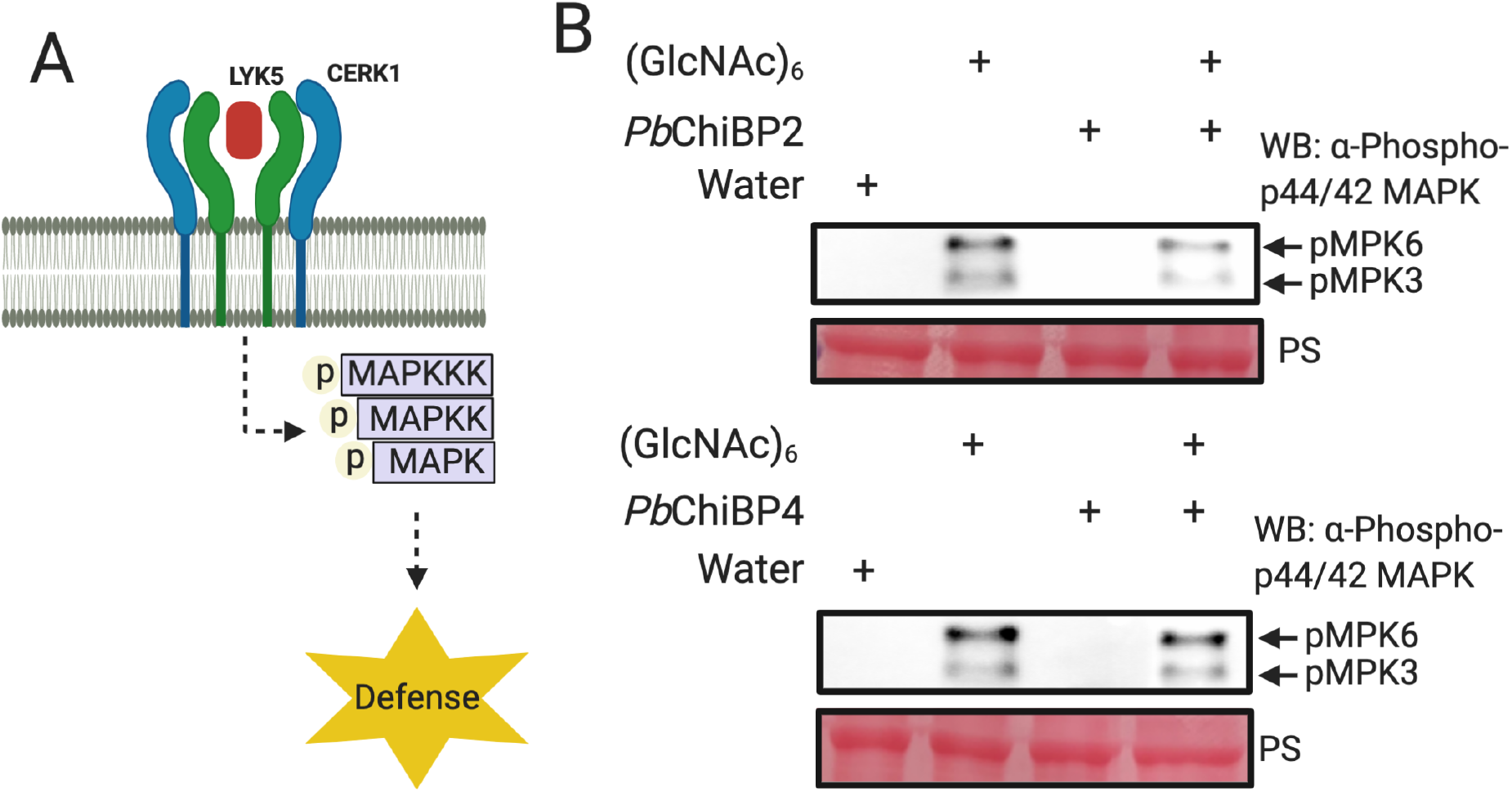
*Plasmodiophora brassicae Pb*ChB2 suppress chitin-induced activation of *Brassica napus* immunity-related mitogen-activated protein kinases MAPKs. **A**. Schematic representation of MAPKs activation through chitin-triggered immunity after chitin recognition by plant receptors. **B**. Western blot (WB) analysis of *B. napus* seedlings after treatment with 1 μM chitin (GlcNAc)6 with or without 10 μM *Pb*ChB2 or *Pb*ChB4. PS stand for ponceau stained blots showing the RuBisCO large subunit as protein loading control. Control were treated with water only (mock), *Pb*ChB2 only or *Pb*ChB4 only. Created with BioRender.com.

## DISCUSSION

Historically, chitin-binding effectors have been widely characterized and discovered in fungal plant pathogens (Sánchez-Vallet et al. 2013). We present here the first chitin-binding effectors from a protist, biotrophic soil-borne plant pathogen, the clubroot pathogen. The Brassicas-*Plasmodiophora brassicae* pathosystem has proven to be very challenging, to the extent that since the discovery of the clubroot disease in the 1700s to this day, only two to four effector proteins have been characterized (Feng et al. 2010; Ludwig-Müller et al. 2015; Yu et al. 2019; Pérez-López et al. 2021). Building on the previous report of CBM18 proteins (*Pb*ChiBs)-encoding genes in *P. brassicae* genome (Schwelm et al. 2015), we focused on those predicted to be secreted and apoplastic. In this study we found that all three putative apoplastic *Pb*ChiB proteins (*Pb*ChiB1, *Pb*ChiB2, and *Pb*ChiB4) have a functional signal peptide, suggesting that they might be indeed secreted to the apoplast of the infected plant cell during the infection.

In this study, we found several CBM18 putative secreted proteins, indicating a possible role of those proteins as effectors. Several of the CBM18 domains found in CE4 proteins might promote efficient substrate deacetylation and transformation to chitosan (Sánchez-Vallet et al. 2015). However, until the activity of the CE4 domain is confirmed, we cannot ascertain the specific role of these proteins. Although we cannot discard the hypothesis that several of them could be strictly involved in pathogen cell wall synthesis and development, several of the families reported here include plant pathogen effectors. For example, *Mp*Chi, a *Moniliophthora perniciosa* effector belonging to the GH18 family, lost its chitinase activity over time, but kept its chitin-binding ability preventing chitin-triggered immunity during the infection (Fiorin et al. 2018), or the *Verticillium dahliae* polysaccharide deacetylase *VdPDA1*, an effector that increases the pathogen virulence through the deacetylation of chitin oligomers (Gao et al. 2019).

Here we showed that the expression of the putative secreted chitin-CAZymes was generally host- and life-stage-dependent, evidencing the adaptability of the pathogen and the sole of these proteins during specific life-stages. Similar was observed for *Pb*ChiB2 and *Pb*ChiB4, which transcript was over-expressed during primary infection (*Pb*ChiB4), and during secondary infection (*Pb*ChiB2 and *Pb*ChiB4), more specifically during resting spore development after penetration and during resting spore formation. Taking into account the uniqueness of *P. brassicae* as a plant pathogen, is hard to draw comparison with the expression profile previously reported for other chitin-binding effectors, although something consistent in other studies is the fact that these effectors are overexpressed during biotrophic phases of the infection (Takahara et al. 2016, Volk et al. 2019).

Our results showed that *P. brassicae* effectors *Pb*ChiB2 and *Pb*ChiB4 were able to bind chitin *in vitro* and to suppress chitin-binding immunity. Similar observations were reported for the *V. nonalfalfae* effector *VnaChtBP* during the parasitic life stages of *V. nonalfalfae* and (Volk et al. 2019), In addition, *VnaChtBP* was able to prevent hydrolysis of fungal cell walls against plant chitinases, which might explain why the interaction of *Pb*ChiB2 and *Pb*ChiB4 with resting spores was not suppressed by the addition of clubroot infected root xylem sap. Functionally speaking, it seems that the three chitin-binding domains identified in plant pathogens (CBM14, CBM18 and CBM50), have evolved independently towards a similar role in plants (Volk et al. 2019). Unlike *VnaChtBP*, which has six CBM18 modules, *Pb*ChiB2 and *Pb*ChiB4 only have two, similar to the LysM chitin binding effectors *ChELP1 and ChELP2* from the anthracnose fungus, *C. higginsianum* (Takahara et al. 2016). These effectors, although not able to protect fungal hyphae against plant chitinases, supress chitin-binding immunity and are indispensable for appressorial functionality.

Our study highlights the first example of the role of a chitin-binding effector in a plant pathogenic protist, a very understudied evolutionary group. This expands the phenomenon outside of fungal plant pathogens and suggests the evolution of a common strategy among plant pathogens to escape plant immunity. Until very recently, the general belief was that chitin-binding effectors with a single chitin-binding module were not able to suppress chitin-triggered immunity (van Esse et al. 2007; Kohler et al. 2016; Sánchez-Vallet et al. 2013; 2020). The discovery of *Mgx1LysM* has challenged this concept by providing evidence that effectors with a single chitin-binding domain were also capable to compete with plant receptors for chitin and suppress immunity (Tian et al. 2020). These findings should support further studies into the *P. brassicae* chitin-binding effectors *Pb*ChiB1, which has only one CBM18 module.

Our revised infection model of *P. brassicae* presented in Fig. 6, based on the results obtained in this study, suggests that soon after root penetration, during the transition from spore to primary plasmodium, *P. brassicae* undergoes through changes of the cell wall composition leaving it susceptible to plant chitinases and to the detection of chitin by plant receptor and the subsequent chitin-triggered immunity. A similar process occurs during the formation of the resting spores, at a time where *Pb*ChB2 and *Pb*ChB4 are highly transcribed. It is interesting that *Pb*ChB2 seems to play a better role ‘masking’ the spores during the infection, but we believe that the weak suppression of chitin-triggered immunity registered for *Pb*ChB4 might be related with its affinity by chitin, which needs to be further studied. However, we feel confident that we show strong evidence of the important role that *Pb*ChiB2 and *Pb*ChiB4 might be playing ‘guarding’ and ‘masking’ *P. brassicae.*

**Fig. 6.**
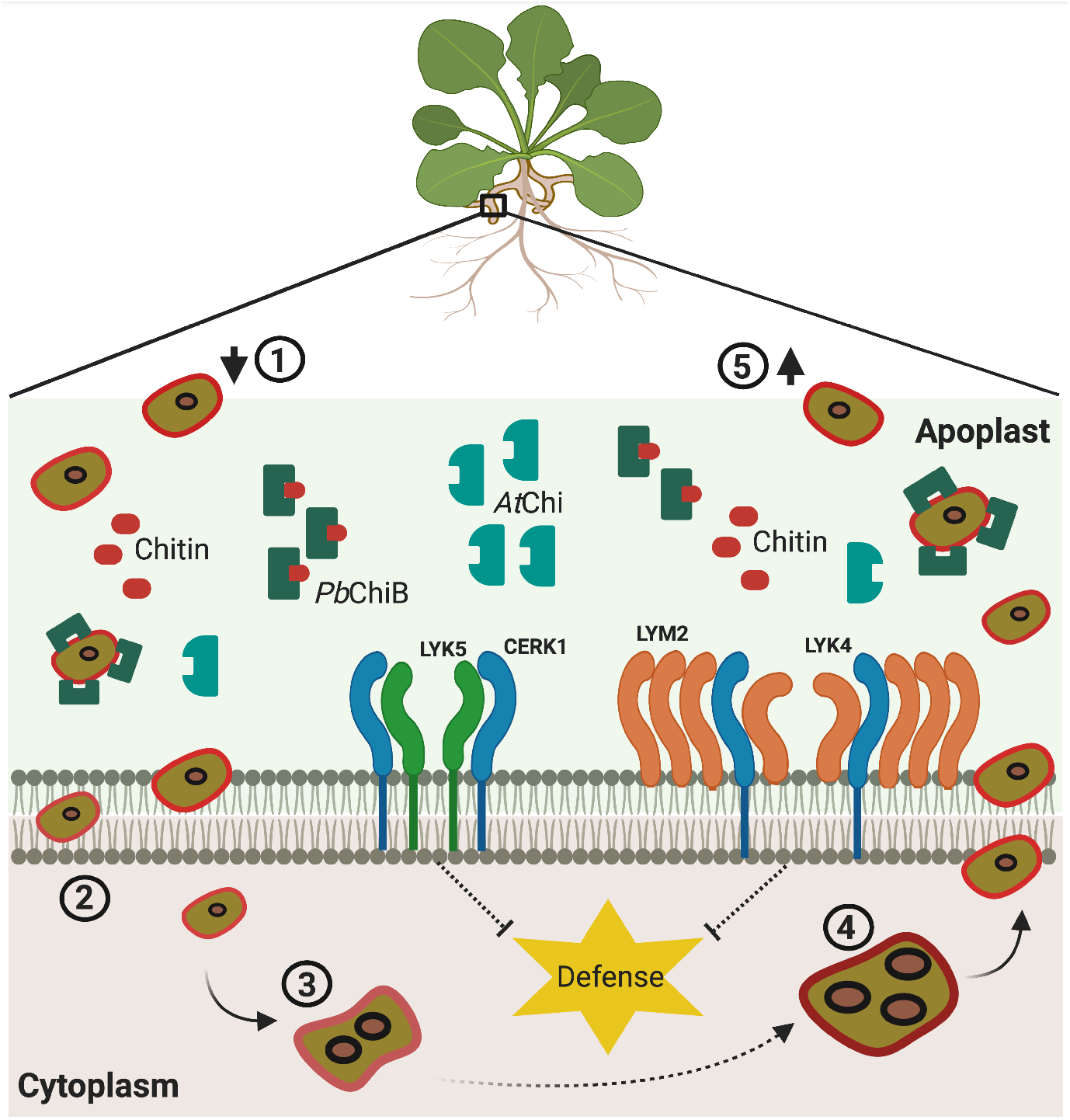
Schematic representation of how the combined activities of *Pb*ChB2 and *Pb*ChB4 guard and mask *P. brassicae* resting spores during infection. Several steps of *P. brassicae* life cycle are presented: (1) Penetration of the roots by resting spores; (2) penetration of the uninucleated primary plasmodium to the cytoplasm of the infected cell; (3) formation of the multinucleated zoosporangia; (4) Formation of the multinucleated secondary plasmodium; (5) exit of the newly formed resting spores from the infected roots. This is a process that could take place several times in the infected plants until eventually the plant dies dehydrated. The possible role of *Pb*ChB2 and *Pb*ChB4 is presenting as a shield of the resting spores and sequestering chitin in the apoplast. Created with BioRender.com.

## AUTHOR CONTRIBUTION

EPL designed the research; KM and EPL performed the research; KM and EPL analyzed the data; KM and EPL wrote and/or edited the manuscript.

## ACKNOWLEDGES

Thanks to Prof. Richard Bélanger and Caroline Labbé from CRIV, Université Laval for the support. Thank you also to Prof. Richard Bélanger and Dr. Tim Dumonceaux for the critical review and comments.

## SUPPLEMENTARY TABLE

**Table S1.**
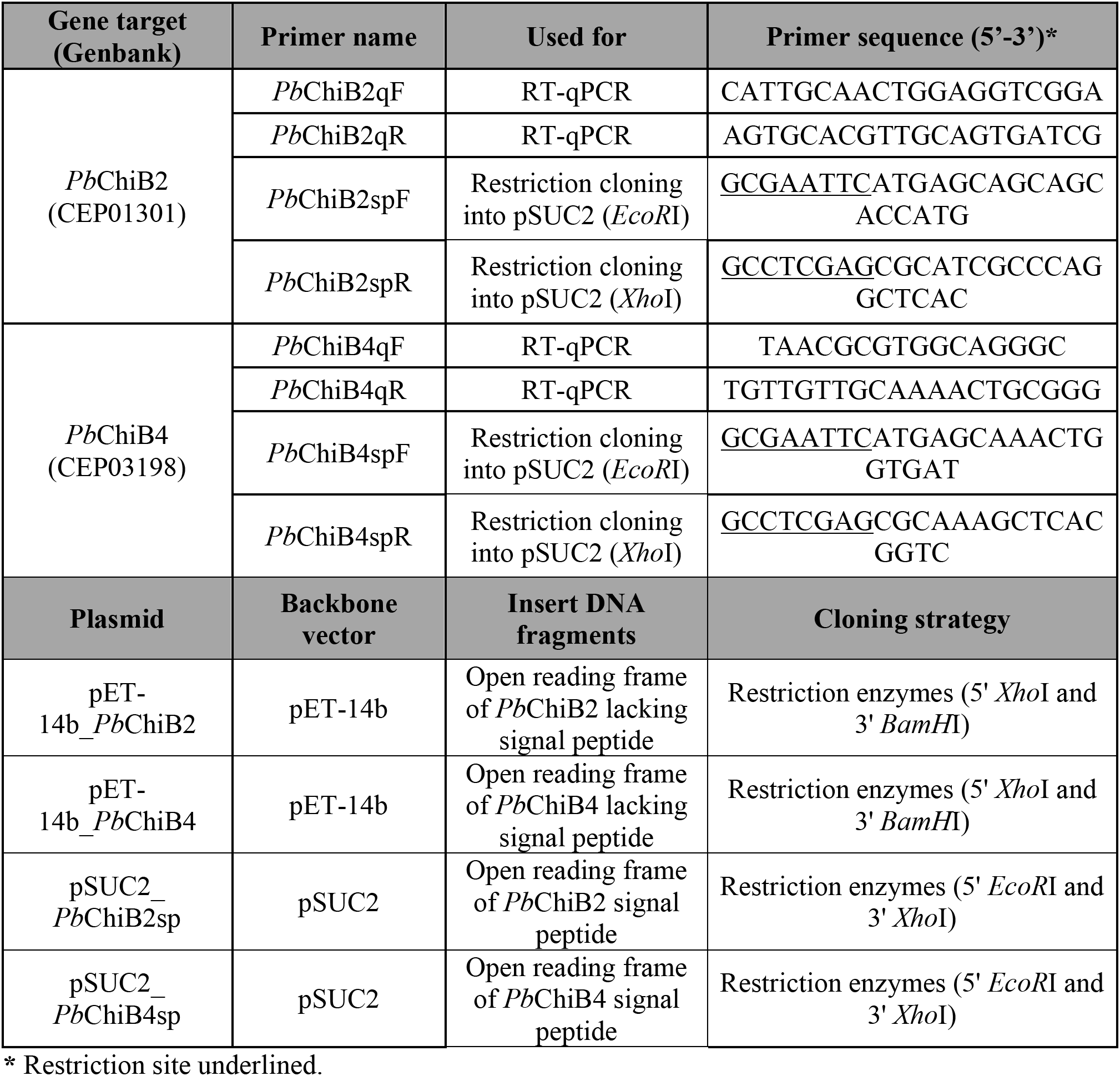
Primers and plasmids used in this study.

## SUPPLEMENTARY FIGURES

**Fig. S1.**
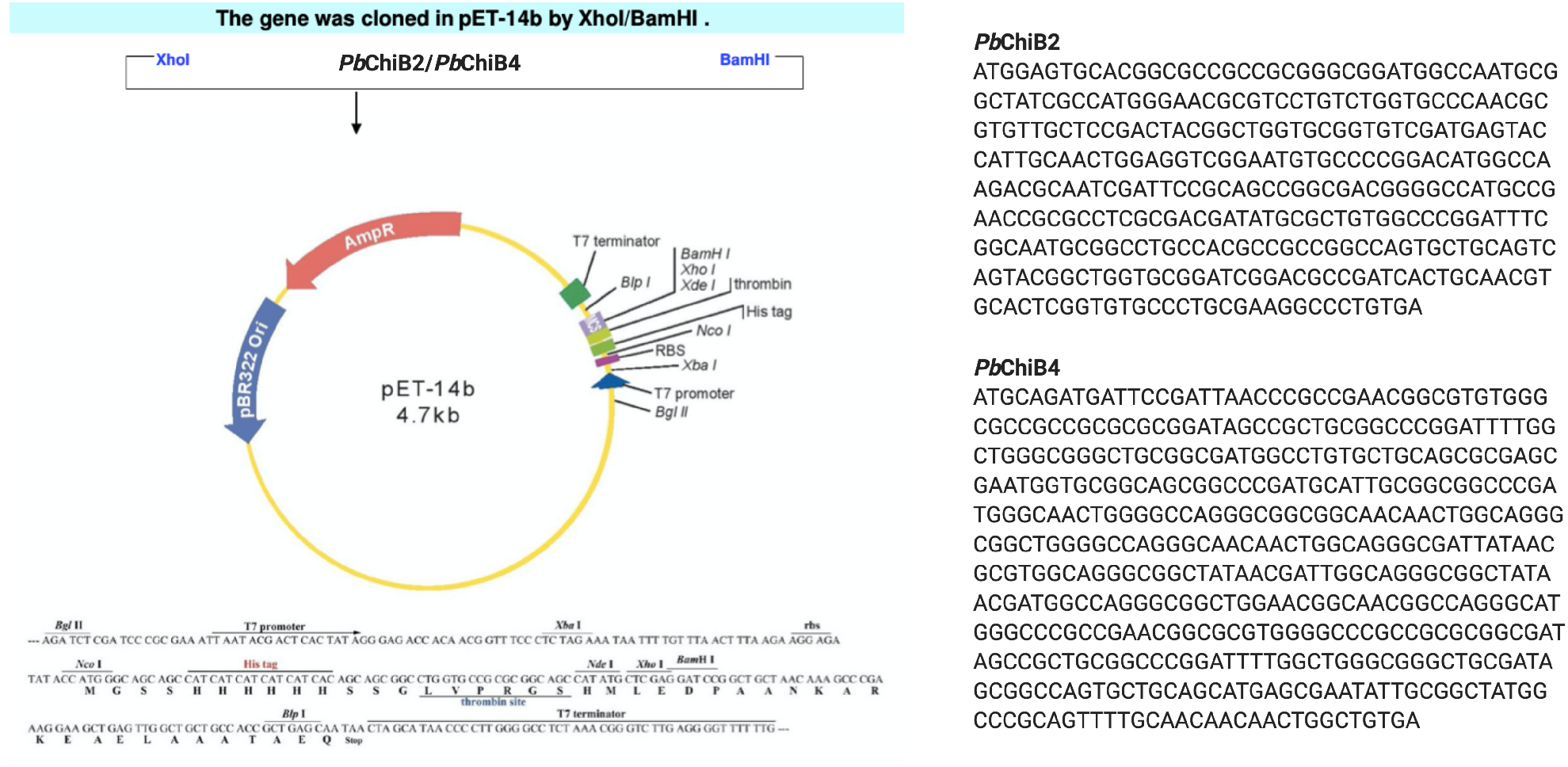
Schematic representation of the cloning strategy. Sequence of *Pb*ChB2 and *Pb*ChB4 encoding genes synthesized by Genscript Biotech. Created with BioRender.com.

**Fig. S2.**
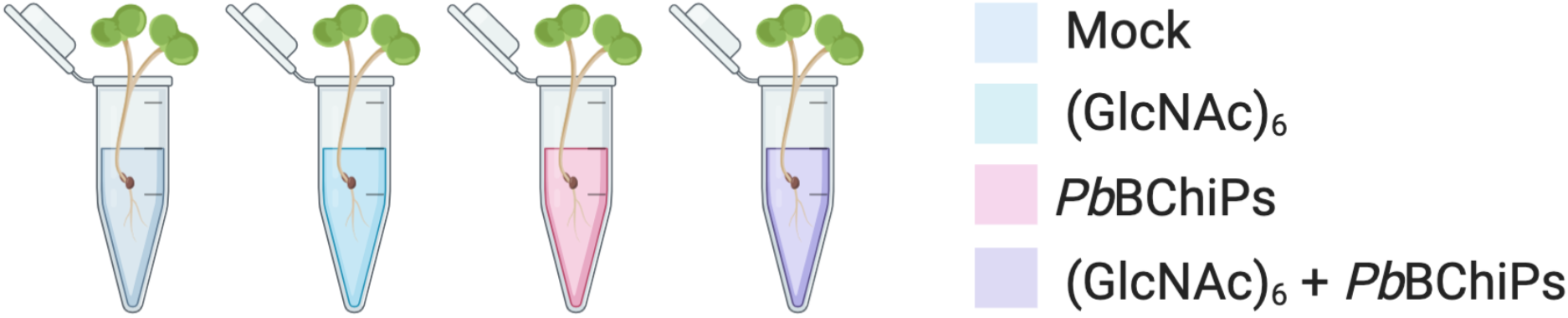
Schematic representation of the MAPK activation assay using *B. napus* seedlings. Three seedlings per treatment were used in each biological replicate of the experiment. Created with BioRender.com.

**Fig. S3.**
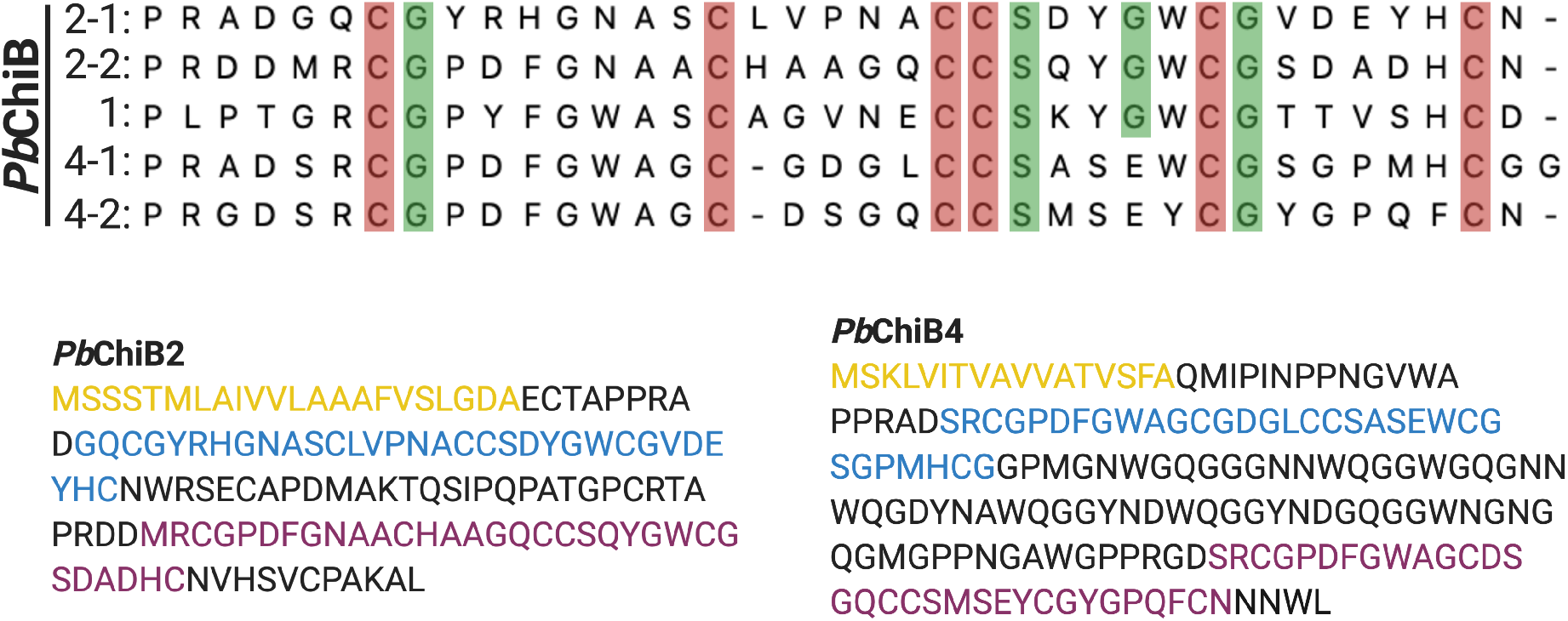
Alignment of *Pb*ChB1, *Pb*ChB2 and *Pb*ChB4 CBM18 modules. In red cysteine residues and in green conserved serine and aromatic residues associated with the hairpin-loop essential for the chitin-binding activity. In the panels the amino acid sequence of *Pb*ChB2 and *Pb*ChB4. In yellow signal peptide (which was removed in the recombinant proteins), in blue the CBM18-1 and in purple the CBM18-2. To see the CBM18 domain structure please check Prosite PS5094 (https://prosite.expasy.org/doc/PS50941). Created with BioRender.com.

